# A Cochlea-Slice Model using Floquet Boundary Conditions shows Global Tuning

**DOI:** 10.1101/2022.10.31.514521

**Authors:** Andrew Tubelli, Hamid Motallebzadeh, John J. Guinan, Sunil Puria

## Abstract

A common assumption about the cochlea is that the local characteristic frequency (CF) is determined by a local resonance of basilar-membrane (BM) stiffness with the mass of the organ-of-Corti (OoC) and entrained fluid. We modeled the cochlea while avoiding such *a priori* assumptions by using a finite-element model of a 20-μm-thick cross-sectional slice of the middle turn of a passive gerbil cochlea. The model had anatomically accurate structural details with physiologically appropriate material properties and interactions between the fluid spaces and solid OoC structures. The longitudinally-facing sides of the slice had a phase difference that mimicked the traveling-wave wavelength at the location of the slice by using Floquet boundary conditions. A paired volume-velocity drive was applied in the scalae at the top and bottom of the slice with the amplitudes adjusted to mimic experimental BM motion. The development of this computationally efficient model with detailed anatomical structures is a key innovation of this work. The resulting OoC motion was greatest in the transverse direction, stereocilia-tip deflections were greatest in the radial direction and longitudinal motion was small in OoC tissue but became large in the sulcus at high frequencies. If the source velocity and wavelength were held constant across frequency, the OoC motion was almost flat across frequency, *i*.*e*., the slice showed no local resonance. A model with the source velocity held constant and the wavelength varied realistically across frequency, produced a low-pass frequency response. These results indicate that tuning in the gerbil middle turn is not produced by a resonance due to local OoC mechanical properties, but rather is produced by the characteristics of the traveling wave, manifested in the driving pressure and wavelength.

**STATEMENT OF SIGNIFICANCE:** The sensory epithelium of hearing, the organ of Corti, is encased in the bone of the fluid-filled cochlea and is difficult to study experimentally. We provide a new method to study the cochlea: making an anatomically-detailed finite-element model of a small transverse slice of the cochlea using Floquet boundary conditions and incorporating global cochlear properties in the slice drive and the wavelength-frequency relationship. The model shows that the slice properties do not show a mechanical resonance and therefore do not produce the frequency-response tuning of the cochlea. Instead, tuning emerges from global cochlear properties carried by the traveling wave.

## INTRODUCTION

The cochlea has a complex, highly ordered anatomy but the role of many of its parts are poorly understood. One way to understand the roles of the various organ of Corti (OoC) anatomical structures and surrounding fluid spaces is to model the cochlea. However, to make a model that is tractable, cochlear anatomy has often been simplified, and/or lumped-parameter representations of parts of the OoC were used. A realistic assessment of OoC motions requires a more realistic model. Of particular interest here is how basilar-membrane (BM) motion is translated into the drives to outer-hair-cell (OHC) and inner-hair-cell (IHC) stereocilia. These bundle motions are currently difficult to measure directly and with a realistic model we can hope to understand these drives. In this paper we present a passive model that is a first step toward addressing the question of how hair bundles are stimulated and thus how cochlear amplification is produced. A future version of this model will include OHC motility to address how cochlear amplification is produced.

To avoid basing our model on pre-formed views of how the cochlea works, we used a finite-element (FE) modeling approach with reasonably accurate anatomy. However, how accurate we can make the material properties of the anatomy is limited by a lack of detailed knowledge about many of the structurally important cochlear elements. FE models can require long computation times, so we included only a 20-μm longitudinal extent of the cochlea in this model, i.e., a cochlear slice. In real cochleae, each cochlear slice is driven by a traveling wave, and to model this we used Floquet boundary conditions on the longitudinal faces of the slice, along with paired fluid-velocity drives at the upper and lower scalae walls with amplitudes set to produce the displacement magnitude and phase of experimental data from the post-mortem gerbil cochlea at the same cochlear region (Meenderink et al., 2022). Such a model configuration allows us to test how global parameters such as wavenumber and frequency interact with local properties such as stiffness and mass, and thus determine if cochlear tuning originates from a local mechanical resonance.

Gerbil was chosen as the model species because gerbils have been used in many cochlear studies, are conveniently small, and have a hearing frequency range that extends to the low frequencies used in human speech (Müller, 1996). Importantly, gerbils are a species on which there are cochlear mechanical measurements using optical coherence tomography (OCT) (e.g., Dong et al., 2018; Fallah et al., 2019; 2021; Meenderink et al. 2022), intra-cochlear pressure measurements (Olson, 1998; Olson, 2001; Kale and Olson, 2015), and measurements of auditory-nerve-fiber responses (e.g., Huang and Olson, 2011; Huet et al., 2016). One disadvantage of gerbils is that their BM is somewhat unusual in that it has an arch-beam architecture (Kapuria et al., 2017). Otherwise, the gerbil has a typical mammalian OoC structure. A preliminary version of this work was presented at the 2022 Mechanics of Hearing Workshop (Tubelli et al., 2022).

## METHODS

### Geometry

The model gross anatomy was based on the Edge et al. (1998) image of the middle turn of the gerbil hemicochlea, which corresponds to a CF of ∼2.5 kHz (Müller, 1996). In a typical cochlear model, the BM is represented as a simple beam, and the upper and lower collagen-fiber layers are considered as a single layer in both the pectinate zone (PZ) and arcuate zone (AZ). In our gerbil model (Fig. 1), the BM was made to have an arched PZ with a smaller arch in the AZ (Xia et al., 2018). The PZ width was two thirds of the total BM width. The PZ upper collagen-fiber layer was a flat shell, and the PZ lower shell layer formed an arch, with ground substance between the shells (Kapuria et al., 2017). The AZ was similar to the PZ but the two fiber layers were closer.

**Figure 1.**
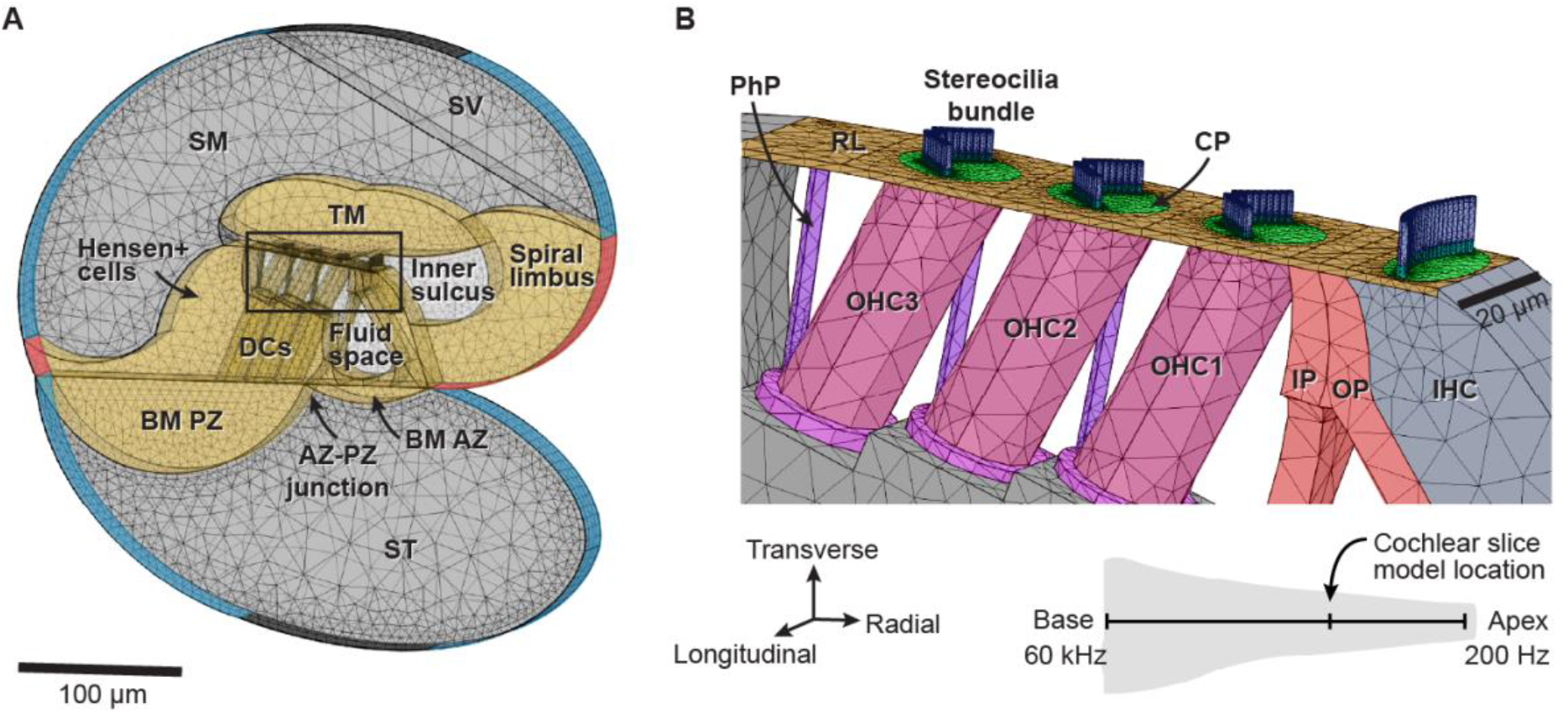
**A**: Slice model geometry with the finite-element mesh, representing the middle turn of the gerbil cochlea. **B**: An enlarged view of the box in A showing anatomical details of the subtectorial space. Bottom right: legend for the orthogonal directions of A and B and the slice location. AZ=arcuate zone; BM=basilar membrane; CP=cuticular plate; DC=Deiters cell; IHC=inner hair cell; IP = inner pillar cell; OHC=outer hair cell; OP=outer pillar cell; PhP=phalangeal process of DC; PZ=pectinate zone; RL=reticular lamina; SM=scala media; SV=scala vestibuli; ST=scala tympani; TM=tectorial membrane.

The slice thickness was 20 μm to make room for a single row of hair cells along with the attachment of the phalangeal processes (PhP) of the Deiters cells (DC) just apical and lateral to the three OHCs, one OHC per longitudinal row (Fig. 1B). Ideally, the PhPs would have been attached to the reticular lamina (RL) three OHCs more apical, but the smaller 20-μm thickness was chosen because it required less computational time. The OHCs, DCs, and PhPs were tilted in the radial direction to mimic the anatomy, but their anatomical tilts in the longitudinal direction were ignored to make them fit within the 20-μm slice.

Within the OoC we included the cells thought to be structurally important (Fig. 1). Structural details were based on gerbil anatomy (Edge et al., 1998; Karavitaki and Mountain, 2007); however, since gerbil data were not available, the diameters of the OHCs, PhPs, and DCs were adopted from mouse data (Soons et al., 2015). The inner pillar cells (IPs) were 20 μm deep and formed a solid wall. The middle parts of the outer pillar cells (OPs) were 10-μm wide with 10-μm gaps that approximate the anatomy.

At the top of the OoC, the RL was a 0.8-μm-thick homogeneous region that included the tops of the pillar cells with inset cuticular plates and PhPs (Fig. 1B). The tops of these structures were all given the same material properties so they effectively formed a single thin plate. The region of the Hensen and Claudius cells lateral to the OHCs and DCs, above the BM, and extending to the spiral ligament, was treated as a single homogeneous region (labeled “Hensen+ cells” in Fig. 1B).

The stereocilia were solid elements, and had fluid-structure interactions like the rest of the model. The tallest stereocilia row was used to model the three rows of stereocilia in a bundle and consisted of a single row of abutting cylinders (Fig. 1B) with appropriate stiffness but no nonlinearity. In order to keep computation manageable, adjacent stereocilia within a row were overlapped so that there were no fluid gaps between them. We omitted the tip links between stereocilium rows and their nonlinearity (OHC motility was not included). The OHC stereocilia formed a single W-shaped bundle with a height (OHC1 and OHC2 were 3.1 μm and OHC3 was 3.0 μm) that spanned the subtectorial gap (i.e., attached at both the cuticular plate and the TM). The IHC stereocilia were 5.3 μm and formed a slightly curved bundle that was attached to the cuticular plate but at the stereocilia tops were freestanding in the fluid. The IHC and OHC stereocilia rows did not span the entire 20-μm slice thickness, which allowed fluid to pass around the bundles (Fig. 1B). The hair-cell and stereocilia dimensions were taken from our own anatomical images of gerbil cochleae. Stereocilia rootlets, which were included in a preliminary version of the slice model (Maftoon et al., 2018), were not included here to reduce the large computational time required for the fine detail of their geometry (Pacentine et al., 2020). The tapering at the base of the stereocilia was omitted to eliminate fluid gaps between stereocilia and therefore reduce computation time. To allow greater bending near the rootlet, we compensated for the lack of stereocilia tapering and rootlets by making the base (lower ∼20%) of each stereocilium bundle a separate material with a ten times lower Young’s modulus.

We combined scala media and scala vestibuli into a single fluid domain that was connected to the fluid in the reticular-lamina (RL) tectorial-membrane (TM) gap and the inner sulcus. Reissner’s membrane is shown for reference in Figure 1A, but the physical membrane was not modeled. There was a second interconnected fluid space inside the OoC (the cortilymph space) composed of the outer tunnel, the fluid surrounding the OHCs, the space of Nuel and (connected via gaps between the outer PCs) the inner tunnel of Corti (Fig. 1A). The fluid domains and soft-tissue regions (i.e., everything except the DCs, hair cells, and stereocilia) extended throughout the 20-μm slice thickness. FreeCAD software was used to create the initial geometry, then coordinates were extracted in order to recreate the shapes for the COMSOL model.

### Material properties

Structural shapes were based on anatomical information available from many sources, but the material properties of most structures have not been measured and had to be estimated. To set initial values and expected limits of values for the Young’s moduli, we used cellular structure (*e*.*g*., the presence and orientation of collagen fibers) and stiffness estimates from the literature. The upper parts of pillar cells have microfilaments and microtubules oriented along a BM-to-RL line (Slepecky, 1996), which implies that this region is relatively stiff, and suggests a relatively high Young’s modulus (Tolomeo and Holley, 1997). The microfilaments and microtubules are randomly oriented in the pillar-cell base (Slepecky, 1996), so the pillar-cell base was modeled as a separate region with an order of magnitude less stiffness. OHC stiffness was estimated based on Tolomeo *et al*. (1996). TM stiffness was based on Sellon *et al*. (2015).

The initial values were then tuned (*i*.*e*., iteratively varied) so that the resulting model motions roughly fit BM radial profile of experimental data (Cooper, 2000; Rhode and Recio, 2000; Homer et al., 2004; Cooper and van der Heijden, 2018) while also keeping the parameter values within estimated physiological ranges and in reasonable relation to each other (e.g., ensuring that the pillar cells were stiffer than the surrounding cells). For the gerbil 2.5-kHz place, there are no experimental data showing the radial profile of BM motion. In the gerbil basal region, the largest BM motion was near the AZ-PZ junction (Fig. 2), and the motion profile was similar for frequencies 12.8-30.4 kHz, and stimulus levels 30–80 dB SPL (Cooper and van der Heijden, 2018). Similar BM radial profiles were found across a range of CF regions in gerbil, guinea pig, and chinchilla (Cooper, 2000; Rhode and Recio, 2000; Homer et al., 2004). Considering this, we assumed that this pattern holds for the gerbil mid-CF region, and we adjusted model parameters using this pattern. The resulting Young’s modulus values used in the slice model are graphically depicted on the model geometry in Figure 3.

**Figure 2.**
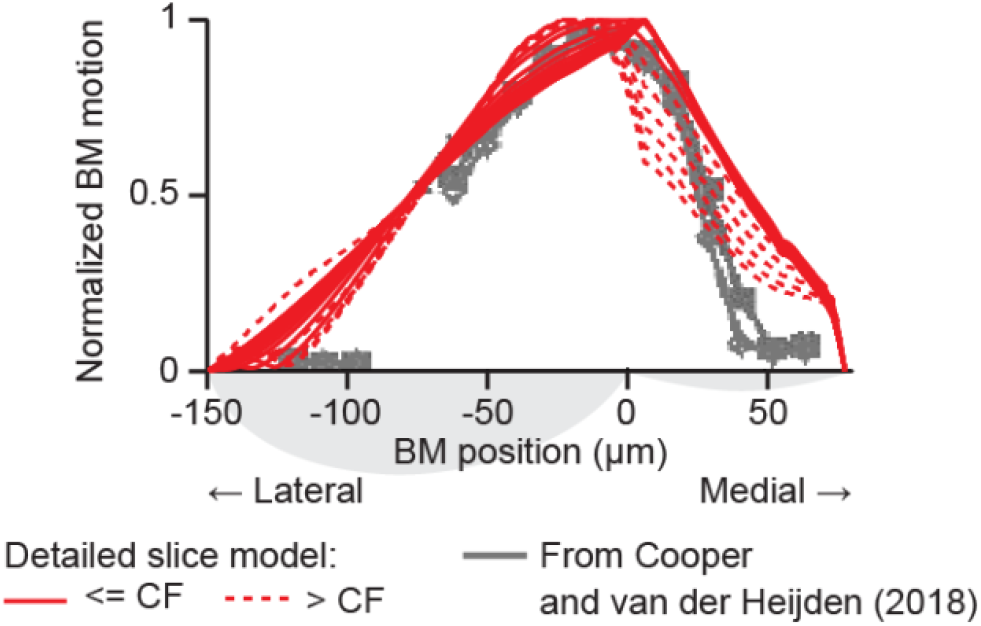
Tuned model basilar-membrane (BM) transverse displacement profile along the radial direction compared to experimental BM motion profiles (Cooper and van der Heijden, 2018). Experimental data are for 12.8 kHz for several sound levels. Red lines show slice model motion from 100 Hz to 5 kHz, solid lines = below CF, dotted lines = CF and above. Each profile was normalized to its peak value. In real cochleae, responses at 4-5 kHz have very low amplitudes and may be where the response is dominated by the fast wave.

**Figure 3.**
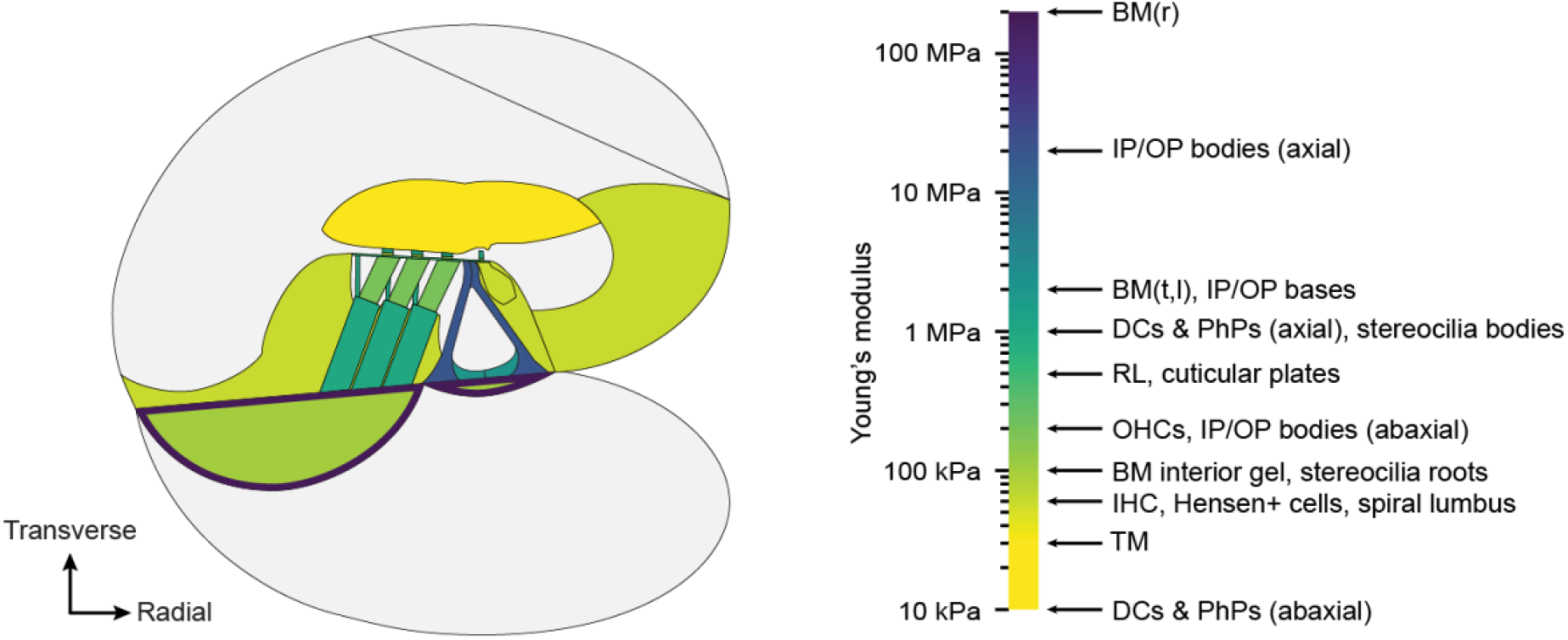
Young’s moduli of model structures with a labeled color scale on the right. For orthotropic materials (BM, pillar cells, DCs, and PhPs), the color denotes the stiffest direction. BM=basilar membrane; IP=inner pillar cell; OP=outer pillar cell; DC=Deiters cell; PhP=phalangeal process; RL=reticular lamina; OHC=outer hair cell; IHC=inner hair cell; TM=tectorial membrane; r=radial; t=transverse; l=longitudinal.

Structural components were elastic and isotropic, except for the BM, DCs, PhPs, and PC bodies which were locally orthotropic. The BM orthotropy followed the collagen-fiber orientation of the upper and lower layers by using a local normal-tangential coordinate system parallel to the outer shell of the BM. The stiffest direction was the tangential direction closest to radial; the longitudinal and transverse directions were two orders of magnitude less stiff based on the more compliant BM ground substance which was a solid with a low Young’s modulus of 100 kPa (Fig. 3), consistent with calculations of ground substance (Naidu and Mountain, 1998). Fiber-layer thicknesses were taken from the ∼2.5-kHz place (Müller, 1996; Schweitzer et al., 1996): the upper (flat) fiber-band thickness was 0.312 μm and the lower (arched) fiber-band thickness was 0.869 μm. The Young’s modulus was 200 MPa for the collagen layers, which corresponds to a volume fraction of 0.2 given a Young’s modulus of 1 GPa for collagen. In each direction, the BM shear modulus was one third of the Young’s modulus, based on the properties of isotropic linear-elastic materials and the assumption that the BM is nearly incompressible. Although the relationship between elastic and shear moduli is more complex for orthotropic materials, we do not have enough information to estimate otherwise. The orthotropic components of the DCs, PhPs, and PC bodies were defined by their local coordinate axes, with the stiffest direction along their lengths and the orthogonal directions two orders of magnitude less stiff. From the model of Motallebzadeh *et al*. (2018), the densities of all soft tissues were set to 1,100 kg/m^3^, the Poisson’s ratios of all solid materials were set to 0.485, and the structural damping was set to 10%. Fluid was considered as water (ten Kate and Kuiper, 1970). We did not model electrical and/or active processes.

### Physics and boundary conditions

The model consisted of three major domains: solid, shell, and fluid, all implemented in COMSOL Multiphysics version 6.0 (COMSOL AB, Stockholm, Sweden). OoC structures were modeled with solid mechanics, except the BM, which was modeled using linear-elastic shell elements with three translational and three rotational displacements comprising the six necessary variables. For the solid and shell components, the momentum equation in its general form is:

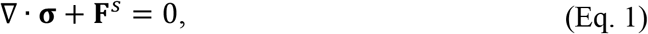

where σ is the stress and Fs is surface traction (gravity effects were neglected). The stress (σ) and strain (ε) relationship for linear-elastic materials, was defined by complex stiffness matrix C, as:

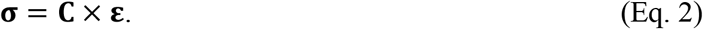

The outermost radial edges of the BM, the medial (toward the modiolus) face of the spiral limbus, and the lateral face of the Hensen+ region were all clamped, i.e., set to zero displacement and no rotation so that the deflection slope was zero.

The fluids were modeled with the thermoviscous-acoustics capabilities of the COMSOL Acoustics module, which used linearized Navier-Stokes equations and included fluid viscosity. Pressure and velocity components in three directions were four variables in the frequency domain. We used the fluid-flow equation of momentum in the frequency domain:

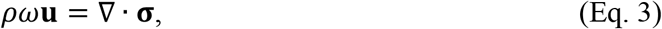

where **u** is the velocity field, **σ** is the stress field (gravity effects were neglected), ω is the angular frequency, and ρ is the fluid density. The stress itself can be expressed in terms of the hydrostatic pressure p and the deviatoric stress tensor τ by

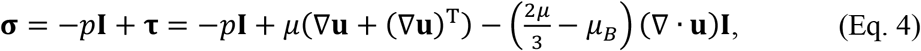

where μ is the dynamic viscosity, μ_B_ is the bulk viscosity, **I** is the identity tensor, and T denotes the transpose. We neglected heat transmission. Rigid, no-slip boundary conditions were applied at the fluid-region walls. At the interface between the viscoacoustic domain and the OoC solid elements, velocity continuity was enforced. At the shared surfaces of shell and solid-shell elements, spatial continuity of displacement was enforced.

#### Wavenumber and the Floquet method

The boundary conditions on the basal and apical faces of the slice need to be specified. In real cochleae there is considerable motion of cochlear structures in the longitudinal direction (Karavitaki and Mountain, 2007; Cooper et al. 2018; Wang et al., 2021), so clamping the slice faces, as was done by Zagadou *et al*. (2020), is not realistic. Instead, we strived to capture the motion induced by the traveling wave by using a computationally-efficient “Floquet” boundary condition that mimicked how the traveling wave changed the phases of the motions at the edges of the slice. At each frequency, a Floquet boundary condition considers the slice to be part of an infinite array of abutted identical slices, each driven identically but at a phase appropriate for a continuous traveling wave. The phase changes are realized by compelling, at each frequency, the phase difference between the basal and apical slice edges to match the phase change of the traveling-wave wavelength being mimicked (Fig. 4). A domain variable of **u** (displacement, velocity, or pressure) at the apical surface (**u**_**2**_) follows the sine-wave periodicity on the basal surface **(u**_**1**_) as a function of wavenumber (**k**) and the respective longitudinal positions z_1_ and z_2_ of those surfaces:

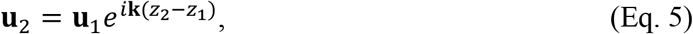

where I is the square root of -1, and e is the exponential Euler number.

In our model, the periodic Floquet boundary condition was applied to the longitudinally facing sides of the model for fluid, solid, and shell domains. In the cochlea, the traveling wave produces an oscillatory pressure on the slice that has a wavelength that depends on the sound frequency and slice location relative to the CF location. As the traveling wave approaches the CF region, its wavelength becomes shorter. Thus, the Floquet wavelength must be changed as a function of frequency to mimic the traveling wave, as is explained in Figure 4. Using a Floquet boundary condition to make a slice model represent the results that would be obtained in a full cochlear model and its traveling wave is a new technique in cochlear modeling that needs to be tested. Such a test is shown in the Supplementary Material and indicates that the Floquet method works.

**Figure 4.**
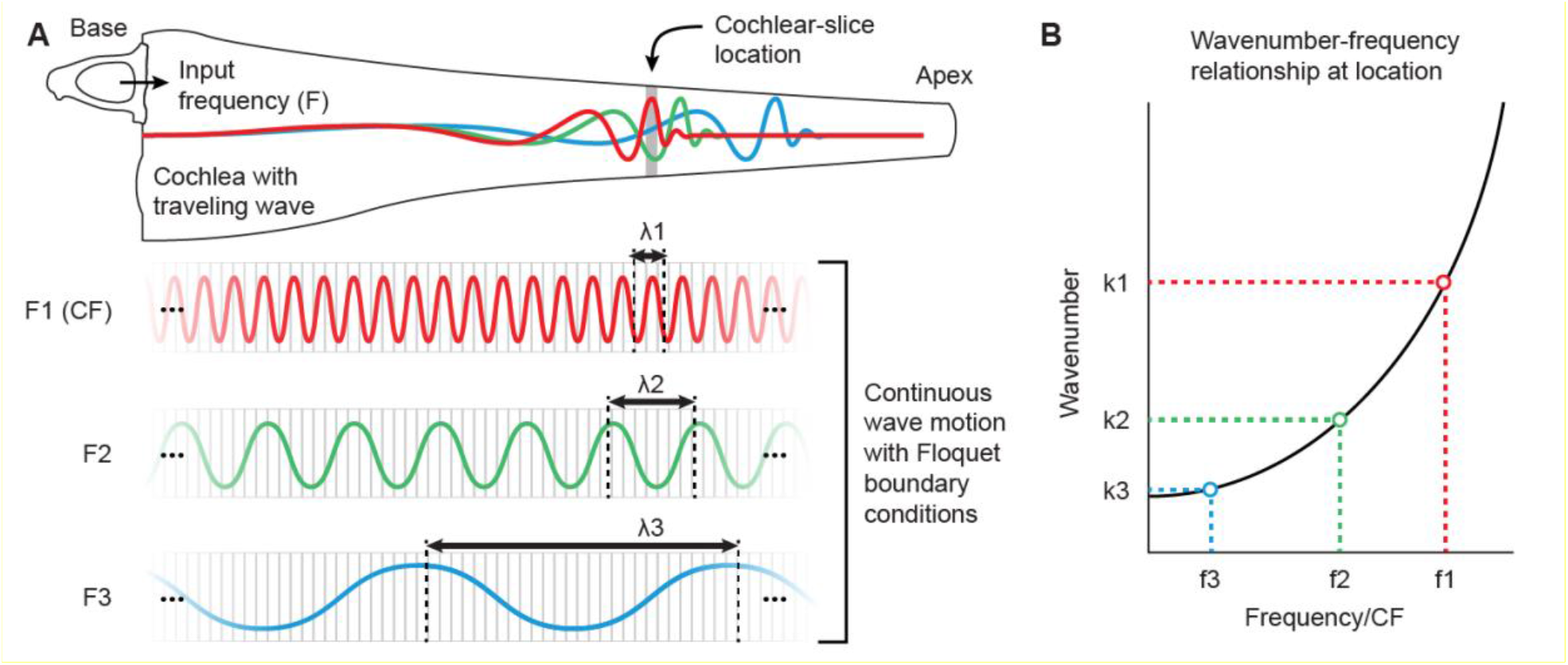
Wavenumber at the cochlear slice related to periodic Floquet boundary conditions. **A**: In the unwrapped cochlear outline at the top, three example BM traveling waves are shown in separate colors for three input frequencies (F). Below are the corresponding sinusoids at the three frequencies overlaid onto an array of slices. Each sinusoid represents the wavelength (λ) of the traveling wave as seen from the slice location. The upper example represents CF with peak motion at the slice location. The other two examples represent lower frequencies with peak motions apical to the slice location and with longer wavelengths at the slice location (λ3>λ2>λ1). **B**: The wavenumber (k), measured in cycles per unit distance, is the parameter specified in the model, and is the reciprocal of the wavelength. In the plot, the solid line is a conjectured wavenumber-versus-frequency relationship (WFR). The datapoint colors correspond to the waveform colors in panel A.

In cochlear models the wavelength is usually represented by its reciprocal, the wavenumber (**k**), which is the number of traveling-wave cycles per unit distance, or spatial frequency. In the cochlea, the wavenumber is a property of the traveling wave that is determined by the gradual changes from base to apex in cochlear dimensions and material properties, e.g., changes in BM width, BM stiffness, and scala areas. For a fixed location, such as that of the slice model or in a typical experiment, the wavenumber is a function of stimulus frequency (Fig. 4). The wavenumber can be complex, with an imaginary part representing decay (or gain) along the cochlea (Shera, 2007), but only the real part representing the spatial frequency is necessary for the slice model.

In the slice model, the wavenumber can be thought of as something that expresses the effects of the rest of the cochlea on the continuous traveling wave as seen at the slice, i.e., the wavenumber vs. frequency relationship (WFR) at the slice is imposed by factors external to the slice. We estimated the wavenumber from the slope of the response phase versus frequency at the 2.5-kHz place of the gerbil (from Fig. 1N of Dong et al., 2018) and the cochlea’s CF-place map (Müller, 1996) using the assumption that measurements across frequency at one place provide information about the response to one frequency across cochlear locations, *i*.*e*., using local scaling symmetry (Zweig, 1976; Sondhi, 1978; Geisler and Cai, 1996). The resulting wavenumber versus stimulus frequency is shown by the thick red line in Figure 5B. WFRs have also been calculated from measurements at multiple locations (Fig. 5A) along the high-frequency region of the gerbil cochlea (van der Heijden and Cooper, 2018) and from a low-frequency region of the gerbil cochlea (Dong et al., 2018); these WFRs are also plotted for comparison in Figure 5B. These alternative WFRs allow us to understand the role of WFR in the model.

**Figure 5.**
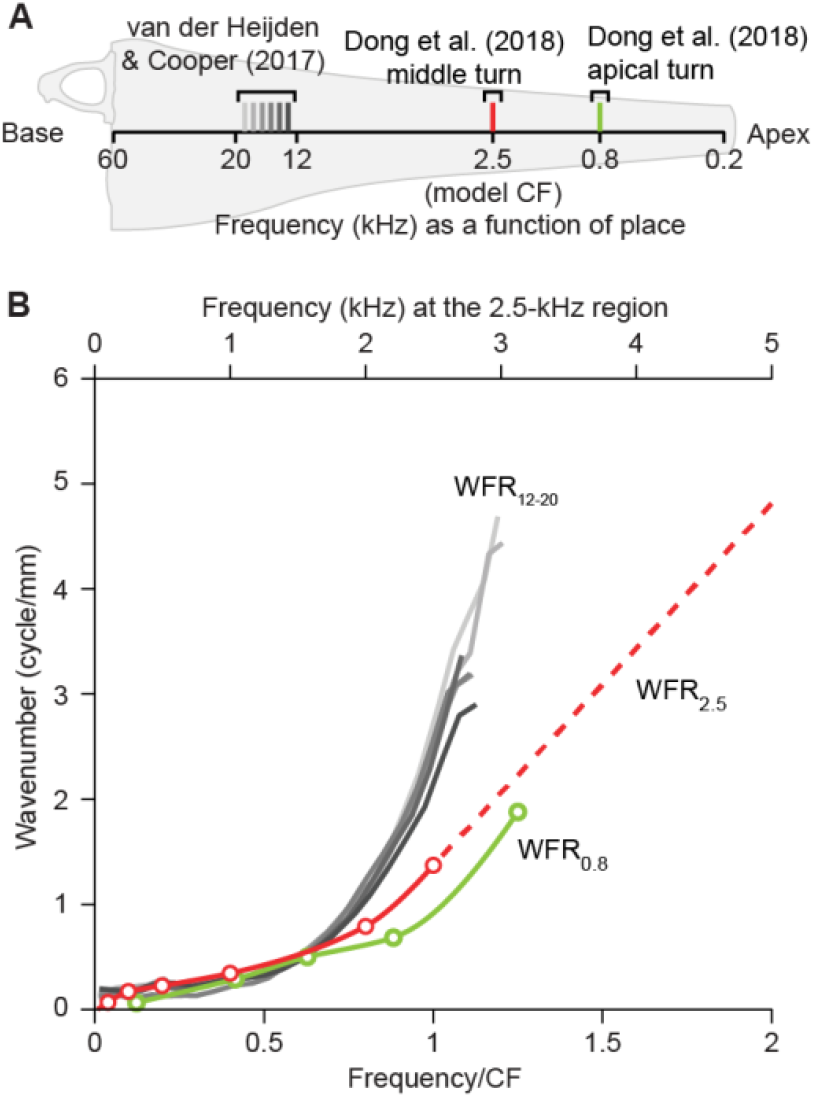
Wavenumber-frequency relationships (WFRs) derived from experimental data. **A**: The locations of the WFR measurements shown in an unrolled gerbil cochlea. **B**: WFR data from the sources and locations in A. The WFR from the 2.5-kHz region (red curves, calculated from Dong et al., 2018) is most relevant for the model. In comparison, the green curve shows the WFR from the 0.8-kHz region, also calculated from Dong et al. (2018) and the gray curves are individual measurements for the ∼16 kHz region, between 12-20 kHz (van der Heijden &Cooper, 2018). All WFRs are normalized to the CF of the measurement preparation. The bottom x axis shows the frequency/CF; the top x axis represents the frequency of the data as translated to the 2.5-kHz slice region. All three WFR solid lines are a cubic interpolation of the data points which, for the estimated WFRs, are shown as open circles. The linear extrapolation at high wave numbers for the 2.5-kHz WFR is represented by the dashed line and was used in the slice model.

#### Model input drive

In real cochleae, sound energy is carried to the slice region by the traveling wave and the resulting scalae pressures that drive the BM vary with sound frequency and cochlear place. There are no measurements of the scala-tympani (ST), scala-vestibuli (SV) or scala-media (SM) sound pressures near the BM in the middle region of the gerbil cochlea. Instead of trying to match unknown scalae pressures, we used sound sources that produced realistic BM motion. The model input was a velocity source pair, one source at the SV top wall and an equal magnitude-and-direction source at the ST bottom wall (input areas can be seen in Fig. 1). Results from a single source at the SV top or at the ST bottom, with a round-window like pressure release at the opposite end, are given in the Supplementary Material. At each frequency, we set input amplitude and phase to mimic the passive BM displacement response from a post-mortem gerbil at a location with a CF close to that of our model (Meenderink et al., 2022; detailed data provided by Wei Dong, personal communication). Since the Meenderink et al. (2022) measurement direction was ∼45° from transverse, the measurements were converted to transverse displacements by multiplying by a √2 (which assumes that BM motion is primarily transverse). The experimental data was measured between ∼0.4-2.4 kHz and did not cover the full range of model frequencies, so at lower and higher frequencies, the experimental data were extrapolated (Fig. 6) using curve fitting to the experimental data (cubic polynomial fit to the log-magnitude, R^2^ = 0.998; and linear polynomial fit to the phase, R^2^ = 0.994). The experimental data also included phase and the phase data was fit by adjusting the phase of the model drive. However, phase is not shown in the figures because the slice model produced only very small phase changes from the driving pressure, so phase plots provide little information.

**Figure 6.**
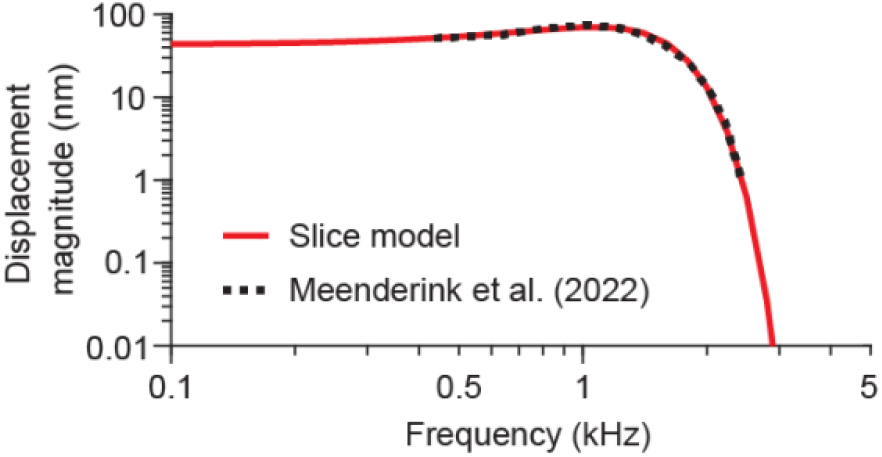
Magnitudes of transverse displacement versus frequency at the BM AZ-PZ junction. The dashed, black line shows the experimental transverse displacement (derived from Meenderink et al., 2022). The solid, red line shows the fitted extrapolated data used in the standard slice model.

### Finite-element computations

Coarse tetrahedral elements were used to mesh the solid and fluid domains, except the viscous layer at the fluid-domain edges used a finer, more structured mesh. The viscous boundary-layer thickness (*δ*_*visc*_) decreased with rising angular frequency (ω) as:

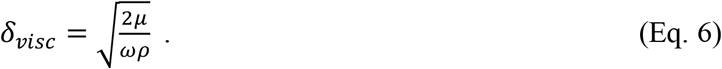

*δ*_*visc*_ was 8.1 μm at 5 kHz, assuming the viscosity (μ) and density (ρ) of water at an ambient pressure of 1 atm and room temperature. The fluid mesh surrounding the OoC and near the bony scalae surfaces were such that there were at least 3 nodes within the viscous boundary layer. Quadratic shape functions were used for the fluid, solid, and shell elements. The number of elements and degrees of freedom were 276,840 and 1,601,146, respectively. A mesh-convergence study was performed to ensure sufficient mesh density for displacement convergence. The simulations were run on Linux on a Dell Tower Station computer with two 20-core Intel® Xeon® Processors for a total of 40 cores (E5-2698 v4) and 512 GB of RAM. Each simulation frequency took approximately 5 minutes.

## RESULTS

The data presented in the main text are for sources at the upper and lower scalae walls (termed the paired-drives model) with amplitudes set to produce the displacement magnitude and phase of experimental data from the post-mortem gerbil cochlea at the same cochlear region (Meenderink et al., 2022). This produces a drive from both scalae and seems most like the normal slow-wave drive. We also made models with the drive just on the upper (SV) wall with a round-window-like membrane on the opposite wall as a pressure release (the SV-drive model), and another model with the drive just on the lower (ST) wall with a pressure release on the opposite wall (the ST-drive model). These models gave slightly different results (see the Supplementary-Material). When the single-source models had equal-amplitude drives, the SV-drive produced more overall OoC motion than the ST-drive. The SV-drive model showed relatively more motion at the OoC top, and the ST-drive model showed relatively more motion at the OoC bottom (see the Supplementary-Material movie). We do not consider or attempt to explain the differences in the motions produced by the different sources. Instead, we focus on model responses that stay qualitatively the same for the different source locations.

### The role of globally determined variables (e.g., wavelength) in the model results

The slice model incorporated anatomical details for the gerbil middle-turn, the ∼2.5 kHz CF region (CF determined by auditory-nerve response maps). However, the slice is part of a cochlea whose dimensions and anatomy change from base to apex. The effects of the more-basal regions on the slice location are carried by the traveling wave to the slice in two ways: (1) by drive amplitude and phase, and (2) by wavelength. The variation of the drive amplitude and phase was accounted for by adjusting the source to reproduce the measured BM motion of Meenderink et al. (2022) (Fig. 7, red line). If the drive amplitude and phase are held constant across frequency (by creating a fixed-drive-amplitude, real WFR model), the result shows much less reduction at frequencies above the local best-frequency (BF) (the BF was ∼1.25 kHz, as derived from the peak of the passive BM motion) (Fig. 7, blue line). The difference between the red and blue lines shows the effects of the drive amplitude falling sharply above BF.

**Figure 7.**
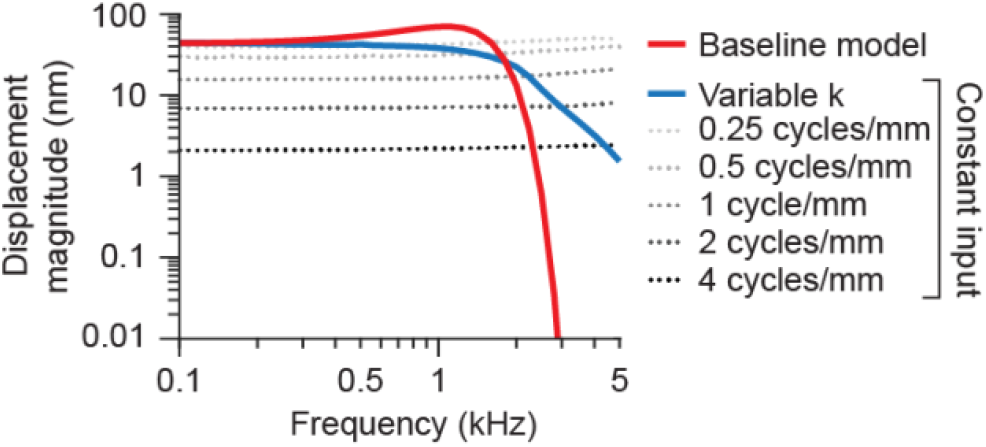
Plots of displacement magnitude for various models and conditions showing the effect of wavenumber on model frequency responses. The red curve is the baseline-model result (shown in Figure 6) for the realistic wavenumber-frequency relationship (WFR) estimated from Dong et al. (2018) and an input set to mimic experimental data (Meenderink et al., 2022). The other curves were from models with a velocity input with a constant magnitude across frequency set to 3 mm/s to align with the low frequencies of the baseline-model curve. The blue curve shows the constant-drive-model response using the frequency-dependent realistic WFR (variable wavenumber, k), and the gray curves show the responses for five wavenumbers held constant across frequency (see key). These results show that the decreasing response of the blue curve at high frequencies comes from the wavenumber becoming higher, thereby yielding lower-amplitude responses. The red curve differs from the blue curve (especially the large drop at high frequencies) because the driving pressure was set to mimic the BM motion, which falls sharply at high frequencies.

The effect on the slice response produced by the variation in wavenumber (or wavelength) with frequency was determined by recalculating the fixed-drive-amplitude model with the WFR replaced by a fixed wavenumber at all frequencies, to yield a fixed-drive-amplitude, fixed-wavenumber model. This was done for five wavenumbers yielding the gray lines in Figure 7. For each wavenumber, the response varied little with frequency but for larger wavenumbers, the response was lower. As shown in the figure, it is the change in the wavenumber as the frequency changes that produces the fixed-drive model’s low-pass responses. These results show that the low-pass characteristics of the fixed-drive-amplitude, real WFR model response are a result of variation in wavenumber rather than variation in frequency response of the local mechanics. In particular, there is no evidence that the frequency response of the slice is shaped by a local resonance due to the local mechanical properties of the slice. Phase was generally affected little by the wavenumber (not shown).

### The overall pattern of OoC deformation

A convenient way to compare relative motions within the slice is to normalize OoC motions relative to the BM motion. For this normalization we used the transverse BM motion at the AZ-PZ junction. The transverse BM motion is the BM motion that has been measured most, so it provides a familiar reference. As expected, the model BM radial and longitudinal motions were much less that BM transverse motion (Fig. 8A-C). Considering this, we normalized the RL and TM motions in all directions by the transverse BM motion (Figs. 8D-I). For this figure, the RL motion was measured as the average along the top surface of the RL, including the three OHC tops. The TM motion was measured on the TM surface across the subtectorial space from the RL.

**Figure 8.**
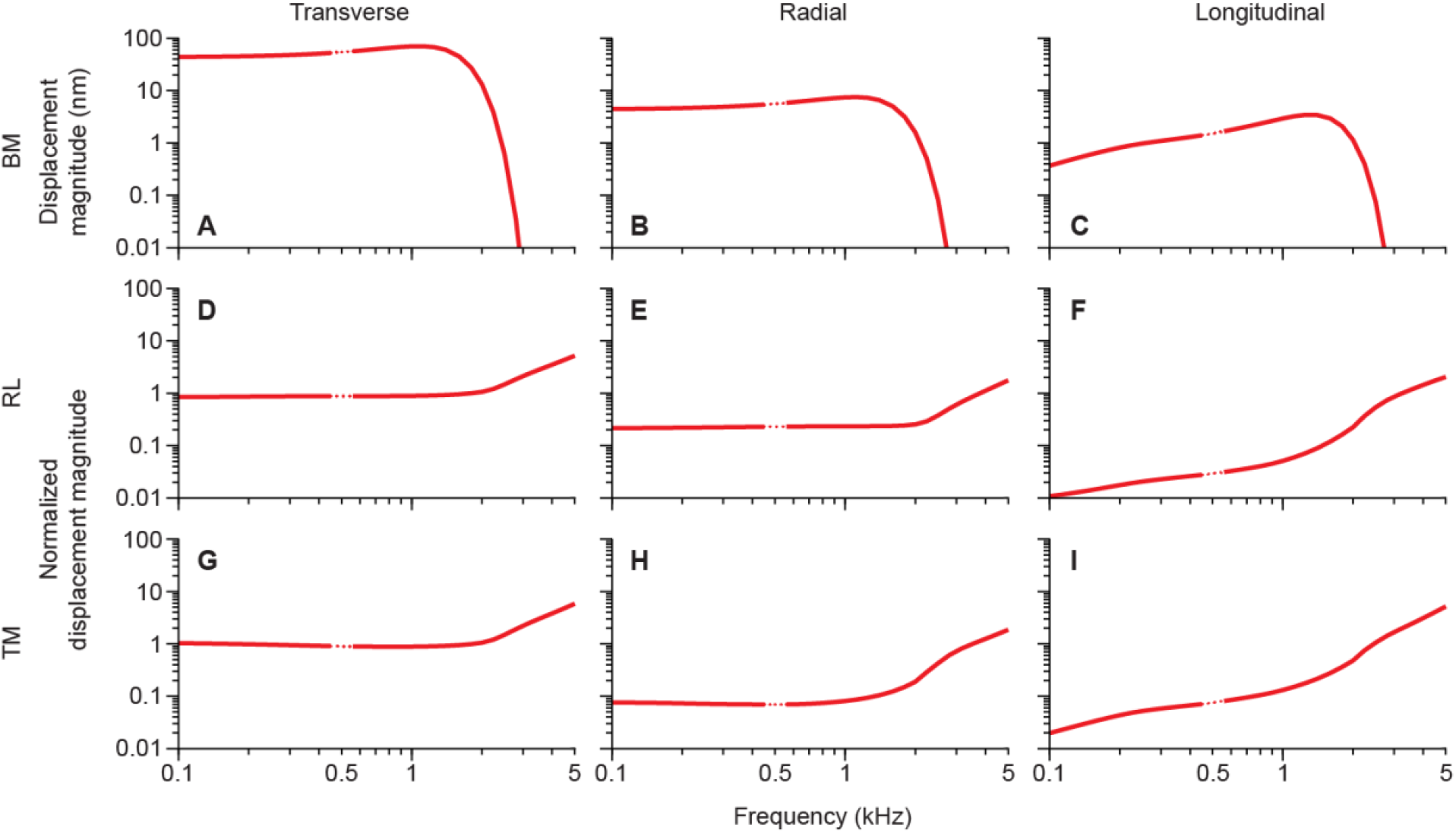
Displacement magnitudes of the basilar membrane (BM) and normalized displacement magnitudes of the reticular lamina (RL) and tectorial membrane (TM). Columns show motions in the directions listed at top. Rows show motions of structures listed at left. The RL (D–F) and TM (G–I) motions in all three directions were normalized by the transverse BM arcuate-pectinate zone (AZ-PZ) junction (A). RL motions were averaged from the tops of the three OHCs, and TM motions were from the TM surface across from the OHCs. The dotted lines at 500 Hz are due to a point that the computation showed to be far different from the nearby points (for an unknown reason), and judging that this was a computational error, we omitted this point.

In Figure 8, the BM transverse displacements have low-pass characteristics, but the normalization removes these characteristics from the normalized TM and RL displacements. The resulting normalized *transverse* displacements are almost flat, with magnitudes ∼1 below CF, which means that these motions follow the BM motion and that all are mostly moving together in the transverse direction (Fig. 8 left column). In contrast, in the *radial* direction the normalized TM and RL displacements, were generally much smaller than the BM transverse motion (Figs. 8E,H) and mostly flat for frequencies below CF. However, this difference produces RL-TM shear (see next section). Finally, the *longitudinal* motions of the TM, RL, and BM are generally much smaller than the BM transverse motion (Fig. 8 right column).

### Relative motions of the drives to the OHC rows and the IHC

It is interesting to look at the differences in motion across the three OHCs and the IHC. Figure 9A-B shows RL and TM radial motion for each of the three OHCs (colored lines), the average of all three (black line—which is typically similar to the response of OHC2), and the IHC. The RL radial magnitudes were substantially different across the OHC rows but the TM radial magnitudes were not (Fig. 9B). Below BF, the IHC radial motion was slightly greater than the motion of the nearby OHC1, in part because it was longer.

**Figure 9.**
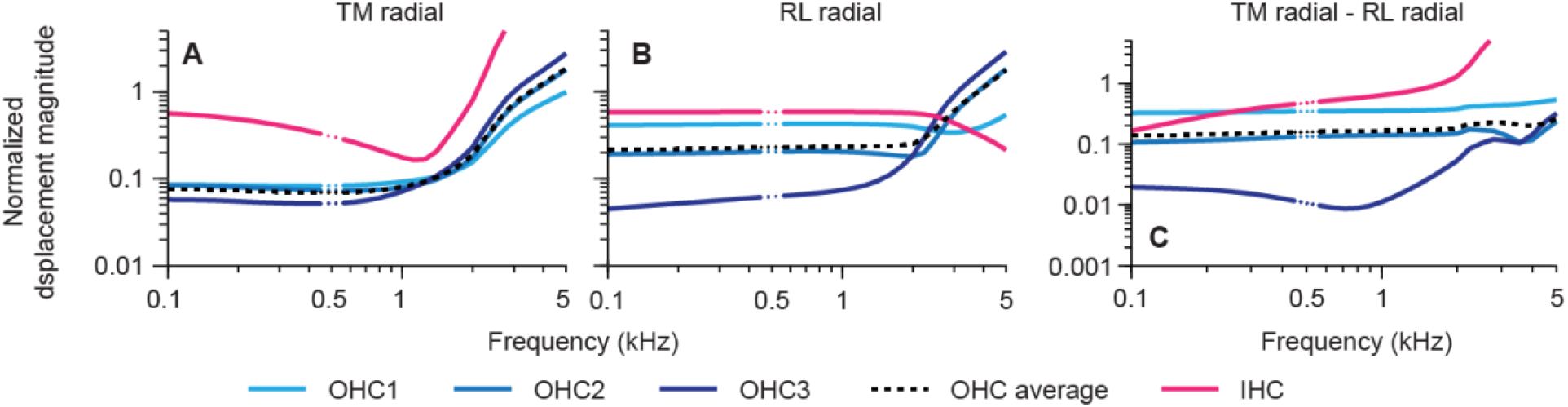
Normalized radial motions of the tectorial membrane (TM) (**A**) and reticular lamina (RL) (**B**) for the three outer hair cells (OHCs) (shades of blue), the average of the OHCs (black), and the IHC (magenta). **C**: The difference (calculated using complex-number values) between the normalized radial motions of the TM and RL. The RL motions were measured by averages along the top surface of each OHC or IHC, and the TM motions were measured as averages at the TM surface directly across the subtectorial gap from the OHCs, or in the case for IHC, at the tops of the stereocilia. All motions were normalized by dividing by the transverse BM motion at the BM arcuate-pectinate zone (AZ-PZ) junction. OHC1 is the OHC closest to the pillar cells and OHC3 is the furthest away. The dotted lines at 500 Hz are due to a point that the computation showed to be far different from the nearby points (for an unknown reason), and judging that this was a computational error, we omitted this point.

The variations in the TM-RL radial motion differences across the three OHC rows are shown in Figure 9C. The difference between the TM and RL motions is of great interest because this difference causes the deflection of the OHC stereocilia that drives OHC motility. The magnitude of the difference between the TM and RL motions was greatest in the radial direction, consistent with the traditional view of the drive to the OHCs. In the important radial motion, OHC1 showed the greatest difference and the motion of OHC3 was much less. The model results indicate that OHC1 receives the largest excitation across the whole frequency range.

### Longitudinal fluid flow

Although the model includes a detailed anatomical representation of only a 20-μm slice, by setting the drive amplitude to match measured BM motion and by using an experimentally-derived WFR, the model incorporates the longitudinal variation of a full cochlea. To explore whether there is longitudinal fluid flow in the model, we chose a point approximately in the center of each of the longitudinal OoC fluid spaces and measured the fluid velocity (there is no structure in these fluid spaces whose displacement can be shown) and for comparison show the velocities of nearby solid structures (Fig. 10). The motions of the four fluid spaces were almost identical in the transverse direction, changing no more than a factor of 2 at the peak. In the radial direction they were similar but the tunnel of Corti (ToC) radial motion was slightly less than the others, presumably because it was closest to the BM and received less radial drive from the OoC rotation. In the longitudinal direction, the space of Nuel (SN) and the outer tunnel (OT) had relatively little motion, presumably because they are narrow spaces and are much-affected by the viscous boundary layers of the surrounding solid structures. In contrast, the greatest motion was in the sulcus, followed by the ToC, consistent with the longitudinal motion being larger when the fluid-space cross-sectional area is larger. Also interesting is that, below the BF region, the radial and transverse motions increase with frequency at approximately 6 dB/octave, however the longitudinal motion increases faster than that, by 10-12 dB/octave. In comparison, the motions of adjacent solid structures (Fig. 10) and are much less in the longitudinal direction.

**Figure 10.**
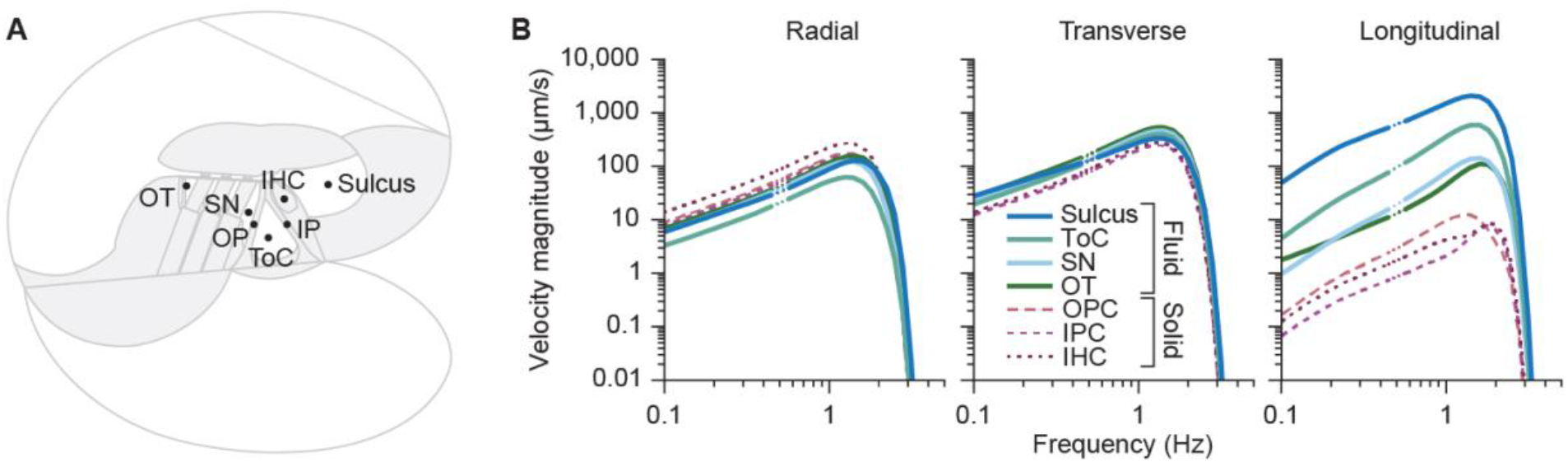
Velocity in the three orthogonal directions measured at a central point in each of the four fluid spaces – the spiral sulcus, the tunnel of Corti (ToC), the space of Nuel (SN), and the outer tunnel (OT) – and three nearby solid structures – the outer pillar cell (OP), inner pillar cell (IP), and inner hair cell (IHC). **A**: A cochlear cross-section showing where the point data were taken. **B**: Velocity magnitude of each data point in the three directions. The dotted lines at 500 Hz are due to a point that the computation showed to be far different from the nearby points (for an unknown reason), and judging that this was a computational error, we omitted this point.

## DISCUSSION

This is the first cochlear model based on a finite-element representation of a cochlear slice that has detailed anatomy and that has boundary conditions set by the Floquet boundary condition method. We tested the Floquet method by applying it to a slice of a published full-length (but very simple) box cochlear model, and found that the Floquet approach worked well when the source pressure was adjusted to produce the differential pressure across the BM found in the box model (Supplementary Material). How does a Floquet model compare with what happens in a real cochlea? The Floquet method sets up an infinite array of slices, each with a source, with the edge and source phases adjusted to mimic those of a traveling wave at a particular frequency. In contrast, the real cochlea is not infinite and the source is the stapes drive with the round window as a pressure release. This produces a fast-wave that results in a slow traveling-wave that is the main BM and OoC drive (Peterson, and Bogert, 1950). The energy of the traveling wave is carried by the fluid pressure and motion (Lighthill, 1981) that then produces BM up and down motion, and what the Floquet method does is approximate the up and down drive produced by the traveling wave in the local area around the slice. Locally, the actual slow-wave exerts an approximately sinusoidally-varying drive, in both time and distance along the cochlea, and its local effect on the slice is what the Floquet method mimics. The local slow wave motion produces a pressure in the fluid. In the Floquet method, the local piece of the traveling wave at the slice forms a traveling wave that is carried along the infinite array. To mimic the normal slow-wave drive to the slice, the motion drive was a pair of velocity sources at the top and bottom of the slice with the source amplitude adjusted to produce the experimentally measured motion of the BM. This configuration was able to mimic the BM motion found in the published full-length box model (Supplementary Material).

A second thing that was mimicked was the variation due to the change in wavelength with frequency (the WFR). The specification of a wavenumber that varied with frequency relative to the slice CF allowed the phase difference on each side of the slice to mimic the frequency-dependent wavelength of a real traveling wave. This provides an opportunity to compare models that have waves with constant wavenumbers (*i*.*e*., constant wavelengths) against models with realistic WFRs, *i*.*e*., wavenumbers that increase as the wave approaches CF. The comparison has allowed us to see what aspects of the motion and its frequency response come from the mechanical properties of the slice and the surrounding fluid alone, and what properties come from the slice being driven by a traveling-wave shaped by the characteristics of the surrounding cochlea; this partial decoupling can be done in a model but not in a real cochlea.

### The model indicates that cochlear tuning does not arise from a local resonance

A key result is that the model shows no evidence that CF is produced by a local resonance, *e*.*g*., a resonance of the BM stiffness and OoC mass (which includes the mass of the entrained fluid). This result is consistent with Békésy’s (1960) measurements of the drained cochlea that showed no significant resonance in the auditory frequency range. When the model was executed using a drive that was constant across frequency and had a fixed wavenumber, the resulting motion vs. frequency profiles were almost flat across frequency. In contrast, when the model was executed with a drive that was constant across frequency and a realistic WFR, the response was low-pass (Figs. 7). This result is in direct opposition to the common assumption that a resonance of local stiffness and entrained mass produce the local CF (*e*.*g*., Helmholtz, 1863; Kolston, 2000; de Boer, 1997).

The means by which the increase in wavenumber leads to tuning are illustrated in Figure 7. When the wavenumber is constant, the response amplitudes are almost flat (Fig. 7, dotted lines). However, with realistic WFRs, as the frequency goes up the wavenumber gets higher, and higher wavenumbers produce lower-amplitude responses (see the progression as frequency increases of the dotted lines in Fig. 7). Thus, as the frequency increases above CF, the decrease in the motion of models with realistic WFRs is due to the model response changing from one constant-wavenumber response to a higher-wavenumber response that has a lower amplitude (Fig. 7). Why a higher wavenumber produces a lower-amplitude response remains to be explained, but the wavenumber (or wavelength) is an important variable in controlling such things as the amount of longitudinal flow of fluid within the cochlea (e.g., Guinan, 2022).

The traveling-wave amplitude increases as it moves toward the CF place, and a modification of the local-resonance hypothesis posits that the increased amplitude increases the damping within the OoC, which causes the traveling wave to be largest (*i*.*e*., produce the CF) before it reaches the cochlear place that has a resonance at the stimulus frequency (Lighthill, 1981). Our models with fixed-wavenumber inputs (Fig. 7, dotted lines) showed small increases in the motion at frequencies above CF and response peaks at a many kHz higher than is plotted in the figures, which is consistent with the cochlea having a resonance at the CF frequency with the resonant place located far apical to the actual CF place (*e*.*g*., by more than an octave on the cochlear CF map). However, a resonance at the CF frequency that is far apical to the actual CF place indicates that it is *not* a local resonance that is producing the CF place (de La Rochefoucauld and Olson, 2007).

Several previous cochlear models have shown that the cochlear CF map can be created by the changes in cochlear dimensions (going apically along the cochlea, the BM becomes wider and thinner) (*e*.*g*., Siebert, 1974; Steele, 1974; Steele and Taber. 1979). These models indicated that cochlear tuning is an emergent property created by the changes in the properties of the traveling wave (*e*.*g*., its wavelength and pressure), not by local cochlear resonances. While the present paper is not the first to suggest that cochlear tuning is not created by local cochlear resonances, it is the first to show that the mechanical properties in a cross section of the cochlea (including the mass of the surrounding entrained fluid) do not produce a local resonance that could account for cochlear tuning.

A major factor that shaped the model amplitude versus frequency functions is the variation of the traveling-wave pressure with frequency. The traveling-wave pressure falls very rapidly at frequencies above BF, as shown by measurements in the base of the gerbil cochlea (Olson, 1998, 2001; Kale and Olson 2015). Both the pattern of pressure changes and the increase in wavenumber at and above CF are properties of the traveling wave. Overall, the model response is consistent with the hypothesis that the properties of the traveling wave produce cochlear tuning, rather than a local resonance of the OoC structures.

Also to be considered is that the cochlea appears to have multiple traveling waves, *e*.*g*., the classic BM traveling wave, a somewhat different traveling wave in the TM, and perhaps an additional traveling wave within the OoC separate from the BM and TM traveling waves (Allen, 1985; Lee et al., 2015; Guinan, 2020). A more complex representation of WFR, or multiple WFRs, could be important for cochlear amplification; we do not explore beyond a single WFR.

One theory that the model contradicts is the hypothesis that there is a resonance in the radial motion of the TM at about ½-octave below the BM CF, and that this resonance is involved in producing cochlear tuning and/or the correct phase of TM movement to trigger cochlear amplification (Gummer et al., 1966; Lukashkin et al., 2010; Sasmal and Grosh, 2019). Figure 8 shows that the motion of the TM follows that of the BM at frequencies below CF with no sign of the supposed resonance.

### Drive to the OHCs

An interesting feature of the model is that it shows differences in *radial* motion across the three rows of OHCs. The motion is greatest at OHC1 and least at OHC3 (Fig. 9). This appears to arise because there is more stretching of the RL around OHC1 than OHC3 (Supplementary Material Movie S1). In contrast, there is little difference in the TM motion across from the three OHC rows. As a result, the deflection of the OHC stereocilia (*i*.*e*., the drive to OHC motility) is greatest on OHC1 and least on OHC3 (Fig. 9C). Differences in the motions of the three rows of OHCs have been reported by Karavitaki and Mountain (2007) for OHC excitation by current. It is also noteworthy that acoustic trauma produces the most damage to the first row of OHCs (Liberman and Dodds, 1984), which is consistent with the model finding that the deflection of the OHC stereocilia is greatest on OHC1.

The finding in our passive model that there is more motion on OHC1 than on OHC3 is in the opposite direction from the changes in transverse motion across the three OHC rows in an active cochlea (Cho and Puria, 2022). Presumably, in the active cochlea forces exerted on the RL create a greater drive to OHC3 than to OHC1, perhaps because of action of the Deiters cell PhP stretching the RL. Both our passive-model data and the active cochlea data indicate that the RL does not act as a rigid plate.

### Longitudinal Fluid flow

This is the first model with sufficient anatomical detail to provide a realistic, credible assessment of longitudinal flow along the OoC fluid spaces. The largest longitudinal motion was found in the sulcus and appears to be due to BM motion producing a rotation of the OoC around the foot of the inner pillar cell so that the IHC region rotates into and squeezes the sulcus. Several previous papers have suggested that there is longitudinal fluid flow in the sulcus from this origin (de Boer, 1993; Steele and Puria, 2005).

Recently, a body of theory was developed that explains traveling wave amplification by changes in OoC area (Altoè et al., 2022; Guinan, 2022) from OHCs cyclically squeezing the OoC transversely and producing fluid flow along the OoC tunnels (Guinan, 2022). Longitudinal flow of fluid in sulcus was not considered but it could also be involved insofar as squeezing the OoC expands it into the sulcus.

The longitudinal fluid flow in the Sulcus was a strong increasing function of frequency. The squeezing of the Sulcus, whether from rotation of the OoC causing motion of the IHC region toward the sulcus, or from OHC contractions squeezing the OoC in the transverse direction in the active case and therefore expanding the OoC in the radial direction, pushes fluid into the sulcus at one location, and ½ wavelength away along the cochlea the OoC is moving in the opposite direction and sucking fluid out of the OoC. The closer these points are, the larger the flow can be expected to be, i.e., the flow should increase as the wavelength gets shorter – which it does as frequency increases. So, this is another aspect of cochlear mechanics where wavelength may play an important role.

## CONCLUSION

We conclude that the stiffness of the tissue in a cross section of the cochlea, along with the mass associated with the cross section, does not produce a resonance that accounts for the cochlear CF map. Instead, the traveling-wave transmits global cochlear properties to each slice along the cochlea and determines cochlear tuning. In particular the traveling-wave motion profile and the local wavenumber-frequency relationship are important in shaping the frequency response of the cochlea. Finally, a slice model with Floquet boundary conditions provides a model platform that can allow examination of the roles of various OoC elements at a much greater detail than previous models.

## AUTHOR CONTRIBUTIONS

S.P. and J.G. conceived and designed the project. A.T. and H.M. created, modified, and analyzed the results of the model and everyone contributed towards the final model creation. All authors contributed to the writing process. S.P. supervised the project.

## ACKNOWLEDGMENTS

We thank Kevin N. O’Connor, Yanli Wang, Heidi Nakajima, and Charles Steele, for reading draft versions and providing valuable feedback and editing assistance. This work was supported in part by Grant R01 DC07910 from the National Institute on Deafness and Other Communication Disorders (NIDCD) of the NIH, and the Amelia Peabody Charitable Fund.

