## Supplementary material for "A Cochlea-Slice Model using Floquet Boundary Conditions shows Global Tuning": Supp Material

### Testing the Floquet Method

We tested how well the Floquet boundary conditions mimicked the local traveling wave using a published “full-length box model" of the cochlea (Xia et al., 2018). From this box model we made a 20-µm section “slice” centered 7 mm from the base, termed the “box-slice model”, and applied Floquet boundary conditions on the slice faces as was done in the detailed slice model of the main paper (Fig. S1A, B). From the full-length box model we obtained the wavelength-frequency relationship (WFR) from the displacement phase of the basilar-membrane (BM) arcuate zone (AZ) pectinate zone (PZ) junction (AZ-PZ junction) (Fig. S1-C). For the larger fluid spaces (i.e., the scalae) the WFR of water was used, calculated as 2π*F/c, where F is frequency and c is the speed of sound in water.

The mechanical drive to the box-slice model was a pair of equal sinusoidal velocity sources applied to the top wall of scala vestibuli (SV) and the bottom wall of scala tympani (ST), as was done in the detailed slice model. We set the stimulus velocity sources in the box-slice model so that they produced the BM differential pressure (i.e., from 10 µm above to 10 µm below the BM AZ-PZ junction) as was measured in the full-length box model at 7 mm (Fig. S1-D). The box-slice model was linear so pressure and motion scaled linearly with input velocity. For the 2.5-kHz place, the effects of the fast wave become larger than those of the traveling wave at an octave higher than CF, *i.e.*, at ~4–5 kHz (Huang and Olson, 2011), so the maximum relevant frequency was 5 kHz. The model was solved for frequencies between 100 Hz and 5 kHz in 1/6-octave steps.

The frequency-dependent displacement that resulted from matching the differential pressure of the OoC overlapped that of the full-length model (Fig. S1-E). This shows that by matching two key characteristics, the WFR and the pressure differential, from the full-length model in a slice model with Floquet boundary conditions, we got the same BM displacement result as in the full-length model. Note that since the model is linear, if we had chosen source values based on producing the BM motion found at the 7 mm place of the box model (instead of using the pressure difference across the BM) we would have gotten the same result. When we tried using a wavelength that didn’t vary with frequency (instead of the WFR), the results did not match.

Thus, the Floquet boundary conditions applied in the box-slice model provided a good match to the results of the actual full-length model, which indicates that the Floquet boundary conditions, along with the correct WFR, provide a way of mimicking the effects of the box-model traveling wave as seen in the slice used in the box-slice model.

Floquet boundary conditions are different from the methods used in other models, such as the WKB method (*e.g.*, Steele and Taber, 1979). The WKB method depends on changes along the cochlear length being gradual in relation to the wavelength of the response, and solves for the WFR explicitly. The combined numerical and perturbation methods of Steele (1999) are computationally efficient but require manual handling and iteration. FE modeling with a Floquet boundary condition, while computationally more expensive, removes some of the handling required, and allows us to use fully realistic anatomy.

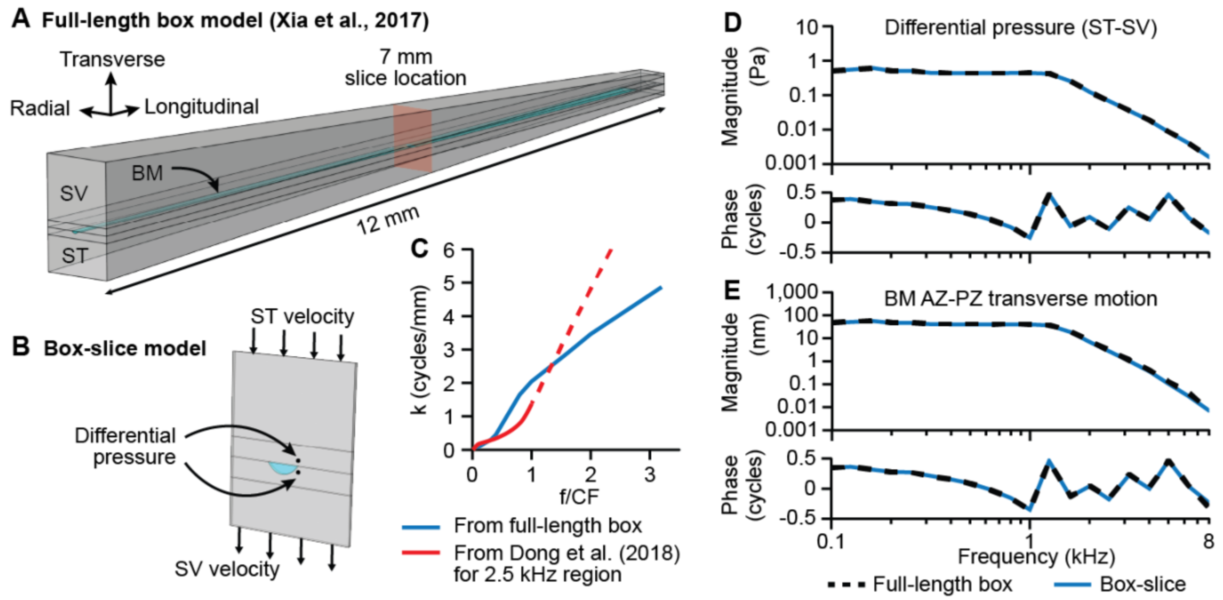


**FIGURE S1**. **A**: Full-length box model from Xia et al. (2018). The SV-ST partition has a BM with an arched pectinate zone but without an organ of Corti. **B**: The 20-µm box-slice model cut from the full-length model. This model was stimulated with a pair of equal (in both magnitude and direction) input velocity sources at the SV and ST walls. Points 10 µm above and below the AZ-PZ zone of the BM indicate where the differential pressure was measured. **C**: The WFR from the full-length model, used in the Floquet boundary conditions for the box-slice model, compared with the WFR from the 2.5-kHz region of the gerbil derived from Dong et al. (2018) that was used in the detailed slice model. The solid regions were estimated from the measurements while the dotted regions are linear extrapolations of the WFRs. **D**: The differential pressure (between scala tympani and scala vestibuli). Dashed lines are from the full-length box model and solid lines show the fitted differential pressure from the box-slice model. **E**: Transverse displacement of the AZ-PZ junction of the full-length model (dashed lines) are matched well by the results from the box-slice model (solid lines). The box-slice model using Floquet boundary conditions produced similar resulting displacements as the full-length model, indicating that Floquet boundary conditions are able to reproduce the response of the full-length model at the slice location.

### Alternative Input Sources

The model geometry was based on the middle turn of the cochlea; however, the scalae pressures that drive the BM are unknown in this region. To get around this, we tuned the source velocity so that resultant displacement was that of experimental data taken at the same region (Meenderink et al., 2022). The main text shows results for a pair of velocity sources: one at the SV top wall and one of equal magnitude and direction at the ST bottom wall. We also tried two alternative single velocity sources for comparison: the ST alone, or the SV alone. In each of these models, the opposite side was given a 10-µm-thick round-window-like shell membrane. The membrane was made to be compliant with a Young's modulus of 100 kPa. Each separate velocity-source model was separately set to produce the experimental data from Meenderink et al. (2022).

The first comparison we made was with BM, RL, and TM motions in the transverse, radial, and longitudinal directions (Fig. S2; like Figure 8 of the main text). There was virtually no difference in the 3 components of the BM displacements (S2, top row) due to stimulation methods. Below the BF of about 2.5 kHz, all normalized RL motions were within a factor of 1.5, and at the higher frequencies ST stimulation produced less RL motion than with ST+SV and SV stimulations. For the TM, ST+SV and SV inputs produced similar TM motions but with a ST input the motion was quite a bit different.

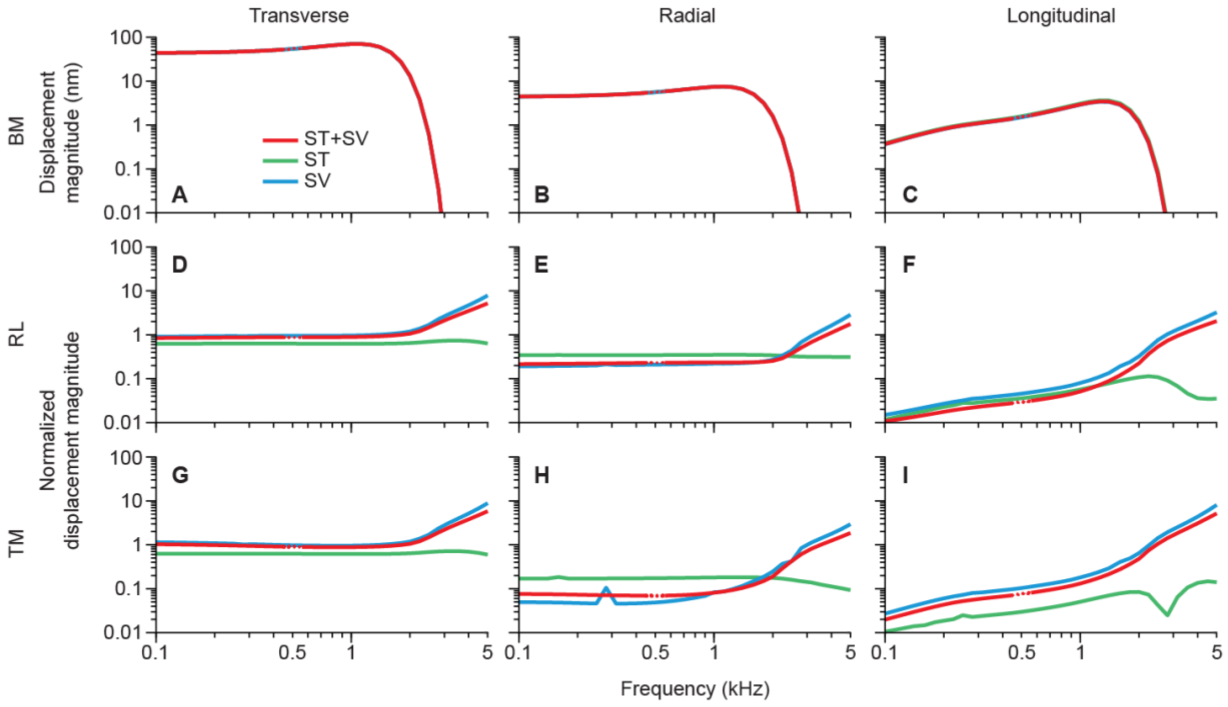


**Figure S2**. Displacement magnitudes of the basilar membrane (BM) and normalized displacement magnitudes of the reticular lamina (RL) and tectorial membrane (TM) for each input combination to the slice model. Columns show motions in the directions listed at top. Rows show motions of structures listed at left. The RL (D–F) and TM (G–I) motions in all three directions were normalized by the transverse BM arcuate-pectinate zone (AZ-PZ) junction (A).

RL motions were averaged from the tops of the three OHCs, and TM motions were from the TM surface across from the OHCs.

We compared the drive to the hair cells (Fig, S3; like Figure 9 of the main text). The results with ST+SV stimulation were more like SV alone than like ST alone. The results with ST were flatter and had less frequency dependance.

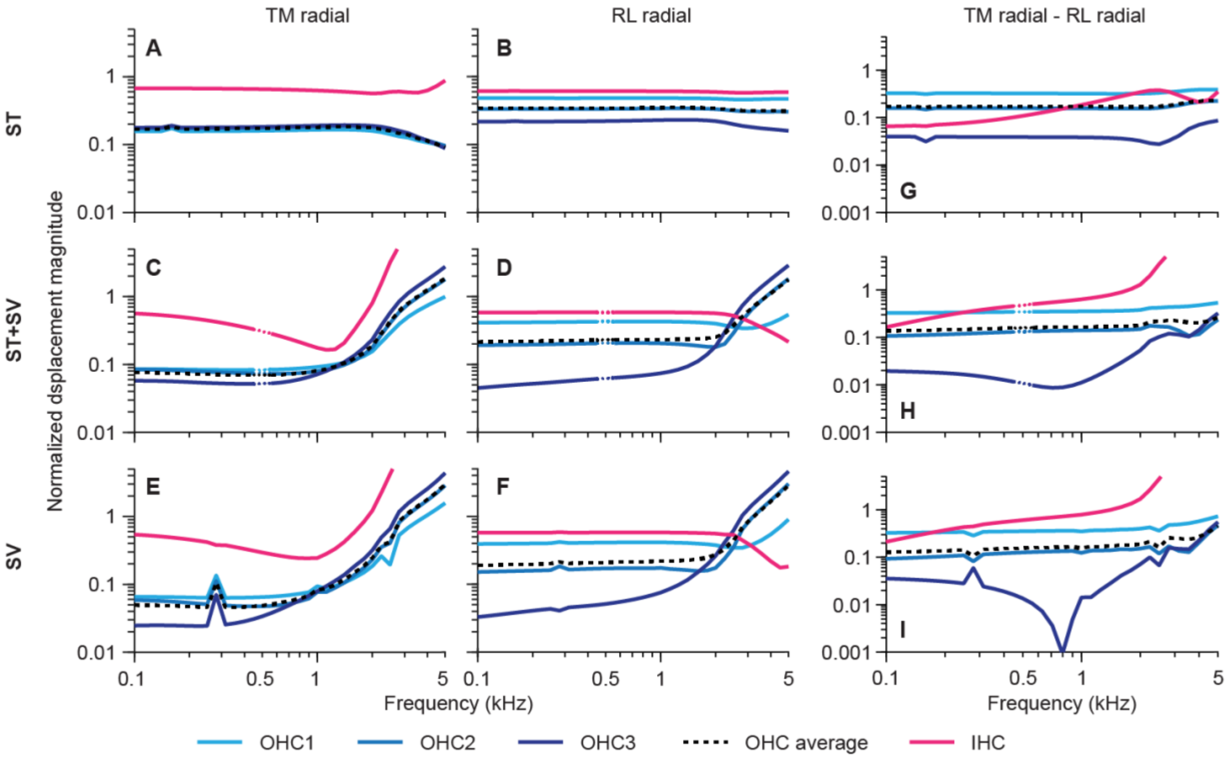


**Figure S3**. Comparison of normalized radial motions for each input combination to the slice model. Normalized radial motion of the tectorial membrane (TM) (A, C, E) and reticular lamina (RL) (B, D, F) for the three outer hair cells (OHCs) (shades of blue), the average of the OHCs (black), and the IHC (magenta). G, H, I: The difference (calculated using complex-number values) between the normalized radial motions of the TM and RL. The RL motions were measured by averages along the top surface of each OHC or IHC, and the TM motions were measured as averages at the TM surface directly across the subtectorial gap from the OHCs, or in the case for IHC, at the tops of the IHC stereocilia. All motions were normalized by dividing by the transverse BM motion at the BM arcuate-pectinate zone (AZ-PZ) junction. OHC1 is the OHC closest to the pillar cells and OHC3 is the furthest away.

### Motion Animations

Animations of motions at a low frequency (400 Hz) and the frequency of peak motion (1.25 kHz) shown in Movie S1 confirm the observations reported above.

**Movie S1**. The overall vector magnitude of the 3D displacement amplitudes of the solid parts of the model, animated over two stimulus cycles for 0.4 and 1.25 kHz, using each of the three velocity source configurations. The superimposed color represents the 3D displacement amplitude. Arrow lengths are logarithmically scaled.
